# Human and Mouse Models of the Bipolar Disorder-Associated *ANK3* p.W1989R Variant Display Altered Somatostatin Interneuron Connectivity and Function

**DOI:** 10.64898/2026.10.23.760718

**Authors:** Kendall P. Dean, Sarah C. Verduci, Clayton C. Kramm, Cooper W. Stinson, Lia Y. Min, Jennifer S. Smith, Cynthia J. DeLong, Mingjia Yao, Kun H. Lee, Jacob M. Reeves, William T. Birdsong, Joshua D. Welch, Paul M. Jenkins

## Abstract

Bipolar disorder (BD) is a debilitating neuropsychiatric disorder characterized by recurrent mood perturbations, cognitive and functional impairment, and increased risk of suicide. Genome-wide association studies (GWAS) have repeatedly identified the gene *ANK3*, which encodes the neuronal scaffolding protein ankyrin-G, as a risk locus for BD. Ankyrin-G interacts directly with the GABA_A_ receptor-associated protein (GABARAP) to stabilize inhibitory synapses, providing a potential mechanistic link between *ANK3* variation and GABAergic dysfunction observed in BD. Previously, we identified an *ANK3* p.W1989R variant in a family with BD and demonstrated that this variant abolishes the ankyrin-G/GABARAP interaction, resulting in inhibitory signaling deficits in mice. Here, we investigated the effects of this variant in human induced pluripotent stem cell (hiPSC)-derived neurons generated from an affected individual heterozygous for the *ANK3* p.W1989R variant. CRISPR correction of the variant increased inhibitory synapse density on both glutamatergic and GABAergic patient-derived neurons. Integrated spatial transcriptomics and calcium imaging identified cell-type-dependent changes in gene expression and calcium dynamics. *ANK3* p.W1989R somatostatin (SST) positive interneurons displayed altered calcium dynamics and reduced SST expression. To determine whether the variant impacts SST interneuron-mediated inhibition of cortical circuits, we selectively stimulated prefrontal cortical SST interneurons in heterozygous *Ank3* p.W1989R mice and recorded optically evoked inhibitory postsynaptic currents (oIPSCs) from cortical pyramidal neurons. SST interneuron-mediated oIPSC peak amplitude, charge transfer, and decay times were reduced in variant mice, without a significant change in the paired-pulse ratio. These findings, using complementary human and mouse models, identify novel roles for ankyrin-G in SST interneuron-mediated inhibitory signaling and demonstrate that altered inhibitory connectivity and SST interneuron function are associated with *ANK3* p.W1989R. This work provides mechanistic insight into how disrupted ankyrin-G-dependent synaptic function may contribute to inhibitory signaling deficits relevant to BD.

## INTRODUCTION

Bipolar disorder (BD) is a chronic and debilitating neuropsychiatric disorder characterized by recurrent episodes of mania or hypomania and depression (1, 2). BD affects more than 1% of the global population and is a leading contributor to disability worldwide (3, 4). In addition to mood instability, affected individuals frequently experience cognitive and functional impairment, reduced quality of life, and elevated rates of medical comorbidities (5-7). BD is also associated with premature mortality and increased risk of suicide, with suicide rates estimated to be 20-30 times higher than in the general population (8). Although the mechanisms underlying BD remain incompletely understood, familial and twin studies estimate BD heritability to be as high as 85%, emphasizing the importance of genetic factors in disease susceptibility (9). Significant progress has been made in BD-associated gene discovery through genome-wide association studies (GWAS), which have identified numerous BD-associated loci. Through these studies, *ANK3* has emerged as one of the most consistently replicated risk genes for BD (10, 11).

*ANK3* encodes ankyrin-G, a scaffolding protein with multiple neuronal isoforms that organizes specialized membrane domains (12). There are three primary ankyrin-G isoforms, which arise due to alternative splicing of exon 37, also known as the “giant exon” (13). Each of these isoforms plays a unique role in mediating cortical neuron excitability and synaptic function (13, 14). The 480 kDa isoform, which contains the entire giant exon, is enriched in the nervous system, and is essential for the organization of excitable membrane domains. The 480 kDa isoform is localized to the axon initial segment (AIS) and nodes of Ranvier, where it clusters ion channels, cell-adhesion molecules, and cytoskeletal proteins required for action potential initiation and propagation (15-17). Beyond these canonical roles, 480 kDa ankyrin-G also contributes to the stability of inhibitory synapses (18, 19). Through these diverse functions, ankyrin-G is an important regulator of neuronal excitability, influencing excitation-inhibition balance within neuronal circuits.

GABAergic inhibitory signaling is essential for regulating neuronal circuit excitability and maintaining proper brain function, and deficits in inhibitory signaling have been associated with several psychiatric disorders, including BD (20-25). Somatostatin (SST)-expressing interneurons represent a diverse class of GABAergic inhibitory neurons that are implicated in cortical plasticity and learning (26). SST interneurons predominantly target pyramidal neuron dendrites, regulating neuronal excitability and network activity through the release of GABA and SST (27). Reduced SST expression has been associated with several mood disorders, with significant reductions in SST mRNA observed in the prefrontal cortex of patients with BD (28, 29). These findings highlight SST-expressing inhibitory neurons as an important neuronal population whose dysfunction may contribute to GABAergic deficits associated with BD.

The role for ankyrin-G in stabilizing inhibitory synapses has been identified in several brain regions, including the cortex, where giant ankyrin-G interacts with the GABA_A_ receptor-associated protein (GABARAP) to prevent GABA_A_ receptor endocytosis and stabilize receptors at the neuronal surface (18). GABARAP is a member of the ubiquitin-like LC3 family of microtubule-associated proteins and interacts with an LC3-interacting region (LIR) encoded within the giant exon of ankyrin-G (19, 30, 31). This interaction requires W1989 within the LIR motif, and substitution of this tryptophan with an arginine (p.W1989R) abolishes high-affinity binding between ankyrin-G and GABARAP (19). Previously, we identified the *ANK3* p.W1989R variant in a family in which all family members diagnosed with BD were heterozygous for the variant (19). In mice, *Ank3* p.W1989R reduces inhibitory synapse density and GABAergic transmission in the forebrain, resulting in pyramidal neuron hyperexcitability and impaired network synchronization (19). However, BD is a genetically complex neuropsychiatric disorder, and it is difficult to create mouse models that truly recapitulate disease phenotypes seen in human patients. Therefore, it is unknown whether the *ANK3* p.W1989R variant will have a significant impact on inhibitory synapse function in a human BD patient genetic background.

The use of human patient-derived induced pluripotent stem cell (hiPSC)-derived neuron models has emerged as a promising technology for the study of genetically complex neuropsychiatric disease, complementary to research being done in animal models (32). Here, we sought to integrate data from human BD patient hiPSC-derived neuronal cultures with a mouse model to better understand the impact of the *ANK3* p.W1989R variant on inhibitory function. We generated hiPSCs from an individual with BD type I who is heterozygous for *ANK3* p.W1989R and used CRISPR correction to generate an isogenic control. Correction of the variant increased inhibitory synapse density on both excitatory and inhibitory hiPSC-derived neurons. Integrated spatial transcriptomics and calcium imaging analyses identified cell-type-dependent changes in gene expression and calcium activity, including reduced SST expression and altered calcium dynamics in SST expressing interneurons with the *ANK3* p.W1989R variant. We then examined SST-mediated cortical inhibition in heterozygous *ANK3* p.W1989R mice using optogenetic stimulation of SST interneurons and identified deficits in inhibitory currents recorded from cortical pyramidal neurons. Collectively, these findings demonstrate reduced inhibitory connectivity and altered SST interneuron function using complementary human and mouse models of the *ANK3* p.W1989R BD-associated variant.

## RESULTS

### Generation of BD patient-derived and CRISPR-corrected isogenic hiPSCs

Previously, we identified the *ANK3* p.W1989R variant in a family in which all genotyped family members diagnosed with BD were heterozygous for the variant (19). Fibroblasts obtained from the male proband with BD type I were reprogrammed into human induced pluripotent stem cells (hiPSCs) through the Heinz C. Prechter Bipolar Research Program at the University of Michigan (Supplemental Figure 1). The *ANK3* p.W1989R variant (c.5965T>C, p.Trp1989Arg) is located within the giant exon of the 480 kDa isoform of ankyrin-G (Figure 1A). To isolate variant-associated effects within the patient’s genetic background, we used CRISPR editing to correct the p.W1989R allele and generate isogenic control hiPSCs (Figure 1B). Sanger sequencing confirmed successful correction of the *ANK3* p.W1989R variant (Figure 1C). The isogenic control retained characteristic hiPSC morphology and expression of the pluripotency markers SSEA-4, SOX2, Oct3/4, and Nanog following CRISPR editing (Figure 1D).

**Figure 1.**
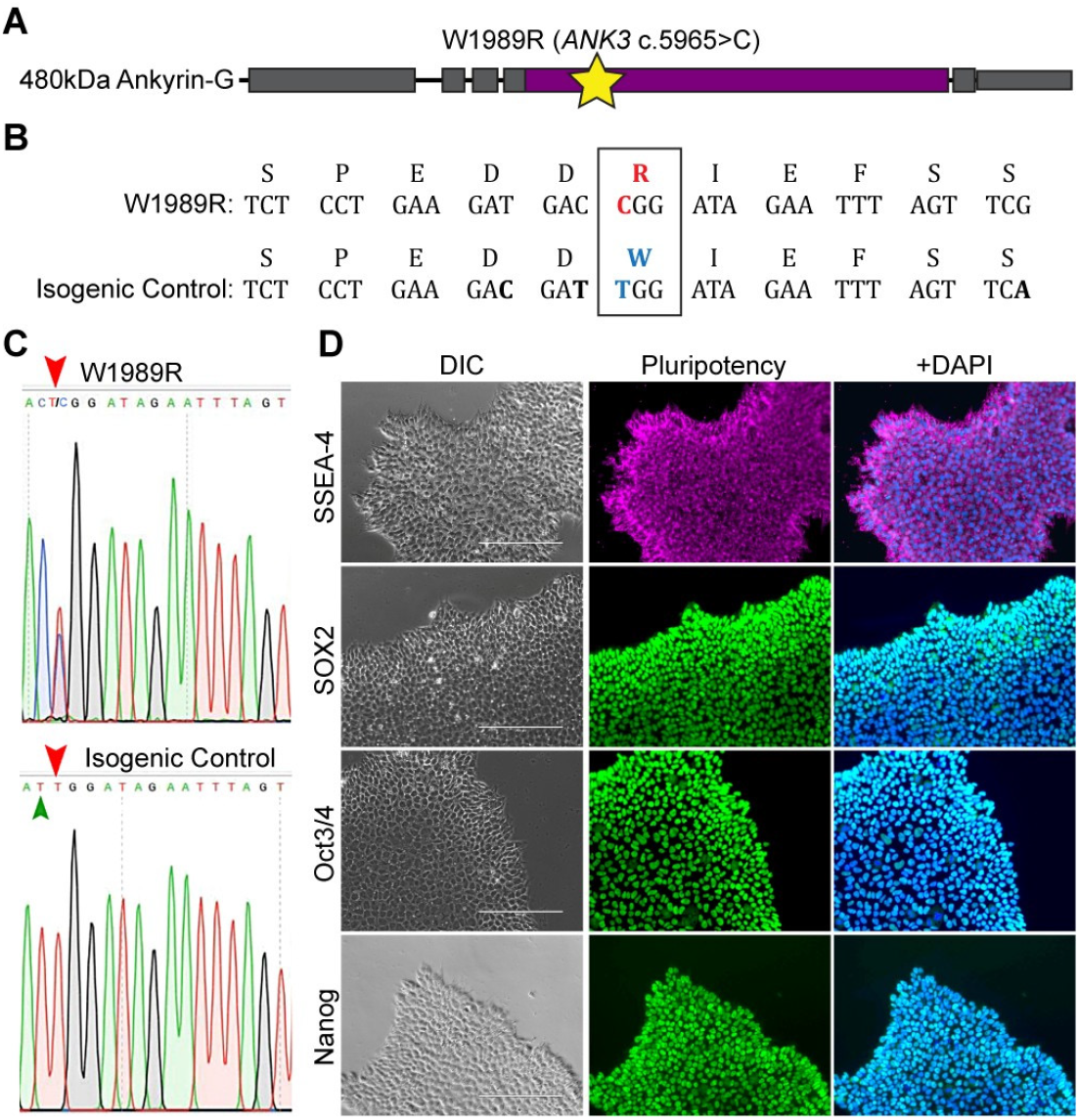
Generation of isogenic control hiPSCs through CRISPR correction of the *ANK3* p.W1989R variant. **A**, Schematic of the 480 kDa isoform of ankyrin-G with the W1989R variant (c.5965T>C, p.Trp1989Arg) marked with a yellow star, located within the giant exon (purple). **B:** Nucleotide coding sequence and single-letter amino acid translation corresponding to the BD patient heterozygous for the non-synonymous W1989R variant (purple), and CRISPR-corrected isogenic control sequence (blue) with inserted silent mutations (bold). **C:** Chromatogram from Sanger sequencing confirming the W1989R heterozygous variant in hiPSCs derived from the affected proband with BD type I (top), and CRISPR correction of the variant and silent mutation insertion to generate isogenic control hiPSCs (bottom). Red arrow: W1989R location. Green arrow: silent mutation. **D:** Differential interference contrast (DIC) microscopy of hiPSC morphology and immunocytochemistry of pluripotency markers SSEA-4, SOX2, Oct3/4, and Nanog. DNA is DAPI-stained in blue. Scale bar: 200 µm.

To generate excitatory and inhibitory neurons from hiPSCs, *ANK3* p.W1989R and isogenic control hiPSCs were engineered using a piggyBac transposable system to insert doxycycline-inducible transcription factors, a nuclear fluorescent marker, and puromycin-resistance cassettes (Supplemental Figure 2A) (36). Doxycycline-inducible NGN2 or ASCL1/ DLX2 constructs were introduced to generate induced glutamatergic (iGlut) or GABAergic (iGABA) neurons from hiPSCs, respectively (Supplemental Figure 2A) (42, 43). Successful piggyBac insertion of transgenes was identified by puromycin resistance and uniform eGFP-NLS or mCherry-NLS nuclear fluorescence. Following 72 hours of doxycycline treatment, RT-qPCR confirmed induction of the corresponding neuronal transcription factors (Supplemental Figure 2B-E).

**Figure 2.**
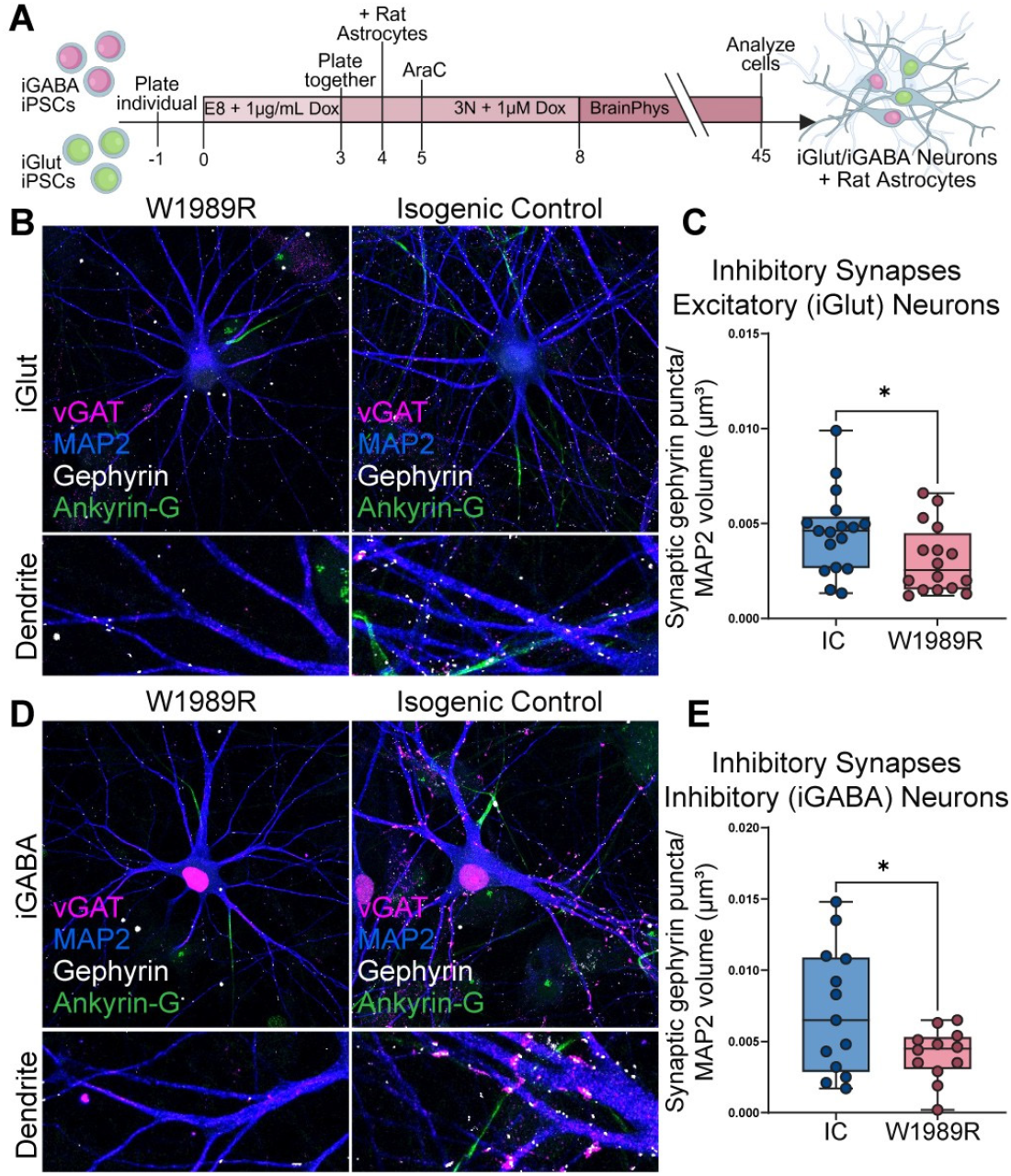
CRISPR correction of *ANK3* p.W1989R increases inhibitory synapse number on iGlut and iGABA neurons. **A:** Schematic for the generation of 45-day-old mixed cultures containing glutamatergic and GABAergic hiPSC-derived neurons (iGlut/iGABA Neurons), co-cultured with rat astrocytes. Cultures were immunostained on day 45 with the somatodendritic marker (MAP2: blue), postsynaptic inhibitory synapse marker (gephyrin: white), presynaptic inhibitory synapse marker (vGAT: magenta), and ankyrin-G (green). Cells were also immunostained for nuclear markers to distinguish glutamatergic (eGFP-NLS: green) and GABAergic (mCherry-NLS: magenta) neuronal types. **B:** Representative images of inhibitory synapses on W1989R and isogenic control iGlut Neurons. **C:** Imaris quantification of the number of gephyrin puncta colocalized with vGAT puncta at the somatodendritic surface, normalized to the volume of the surface, on W1989R (red) and isogenic control (blue) iGlut neurons. **D:** Representative images of inhibitory synapses on W1989R and isogenic control iGABA Neurons. **E:** Imaris quantification of the number of gephyrin puncta colocalized with vGAT puncta at the somatodendritic surface, normalized to the volume of the surface, on W1989R (red) and isogenic control (blue) iGABA neurons. Welch’s t test: *P< 0.05 (iGlut W1989R: n = 16, N = 3, iGlut isogenic control: n = 17, N = 3, iGABA W1989R: n = 12, N = 3, iGABA isogenic control: n = 13, N = 3).

### CRISPR correction of *ANK3* p.W1989R increased inhibitory synapse density on excitatory and inhibitory neurons

Previous work demonstrated reduced inhibitory synapse density on cortical pyramidal neurons in *ANK3* p.W1989R mice, however, it is unknown if these findings would translate to a human BD patient-derived model (19). Additionally, it is unknown if this variant only impacts ankyrin-G-mediated inhibitory synapse stabilization in excitatory neurons, or if inhibitory neurons may also be impacted.

To address these questions, we generated mixed cultures of iGlut and iGABA neurons from *ANK3* p.W1989R and isogenic control hiPSCs and co-cultured them with rat astrocytes for 45 days (Figure 2A). Cultures were immunostained for MAP2, the postsynaptic inhibitory synapse marker gephyrin, the presynaptic inhibitory synapse marker vGAT, and ankyrin-G. Excitatory iGlut and inhibitory iGABA neurons were distinguished by eGFP-NLS or mCherry-NLS labeling, respectively. Inhibitory synapses were identified as gephyrin puncta colocalized with vGAT puncta at the MAP2-defined somatodendritic surface.

CRISPR-corrected iGlut neurons exhibited significantly greater inhibitory synapse density than *ANK3* p.W1989R iGlut neurons (Figure 2B, C). Inhibitory synapse density was also significantly greater on CRISPR-corrected iGABA neurons than on *ANK3* p.W1989R iGABA neurons (Figure 2D, E). These findings demonstrate that correction of heterozygous *ANK3* p.W1989R increases inhibitory synapse density on both excitatory and inhibitory neurons within the BD patient-derived genetic background.

### The AIS began farther from the soma in inhibitory neurons following CRISPR correction of *ANK3* p.W1989R

AIS morphology is dynamically regulated in response to changes in neuronal activity, and previously we demonstrated that *ANK3* p.W1989R mice have decreased AIS length compared to WT counterparts (19, 44). Therefore, we used immunostaining for ankyrin-G to determine if AIS start position or length was altered in *ANK3* p.W1989R hiPSC-derived neurons. In iGlut neurons, AIS start distance and length did not differ significantly between *ANK3* p.W1989R and isogenic control neurons (Figure 3A-C). In iGABA neurons, the AIS began significantly farther from the soma in isogenic control neurons, and AIS length was unchanged (Figure 3D-F).

**Figure 3.**
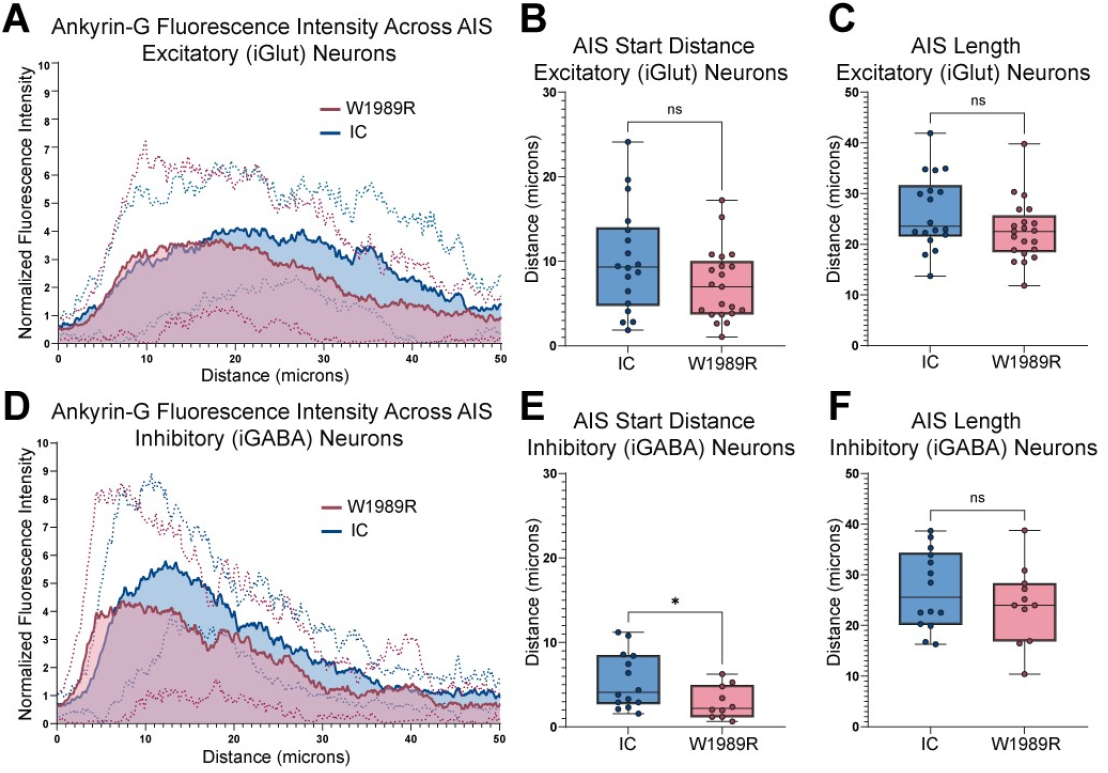
CRISPR correction of *ANK3* p.W1989R increases AIS start distance from the soma on inhibitory (iGABA) neurons. **A:** Plot profile of ankyrin-G fluorescence intensity across the axon initial segment (AIS) for W1989R (red) and isogenic control (blue) iGlut neurons. **B:** AIS start distance from the soma for W1989R (red) and isogenic control (blue) iGlut neurons. **C:** AIS length for W1989R (red) and isogenic control (blue) iGlut neurons. **D:** Plot profile of ankyrin-G fluorescence intensity across the AIS for W1989R (red) and isogenic control (blue) iGABA neurons. **E:** AIS start distance from the soma for W1989R (red) and isogenic control (blue) iGABA neurons. **F:** AIS length for W1989R (red) and isogenic control (blue) iGABA neurons. Welch’s t test: *P < 0.05 (iGlut W1989R: n = 21, N = 3, iGlut isogenic control: n = 18, N = 3, iGABA W1989R: n = 11, N = 3, iGABA isogenic control: n = 14, N = 3).

### VISTA-FISH identified inhibitory and excitatory neuronal populations in *ANK3* p.W1989R and isogenic control using spatial transcriptomics

Video Imaging with Spatial-Temporal Analysis by Fluorescence in Situ Hybridization (VISTA-FISH) combines live calcium imaging with spatial transcriptomics, bridging gene expression profiles and neuronal activity in the same cell at a single-cell resolution (45). To examine the cell-type-specific transcriptional and functional effects of the *ANK3* p.W1989R variant, we performed VISTA-FISH on 45-day-old mixed cultures of iGlut and iGABA neurons from *ANK3* p.W1989R and isogenic control hiPSCs co-cultured with rat astrocytes (Figure 4A).

**Figure 4.**
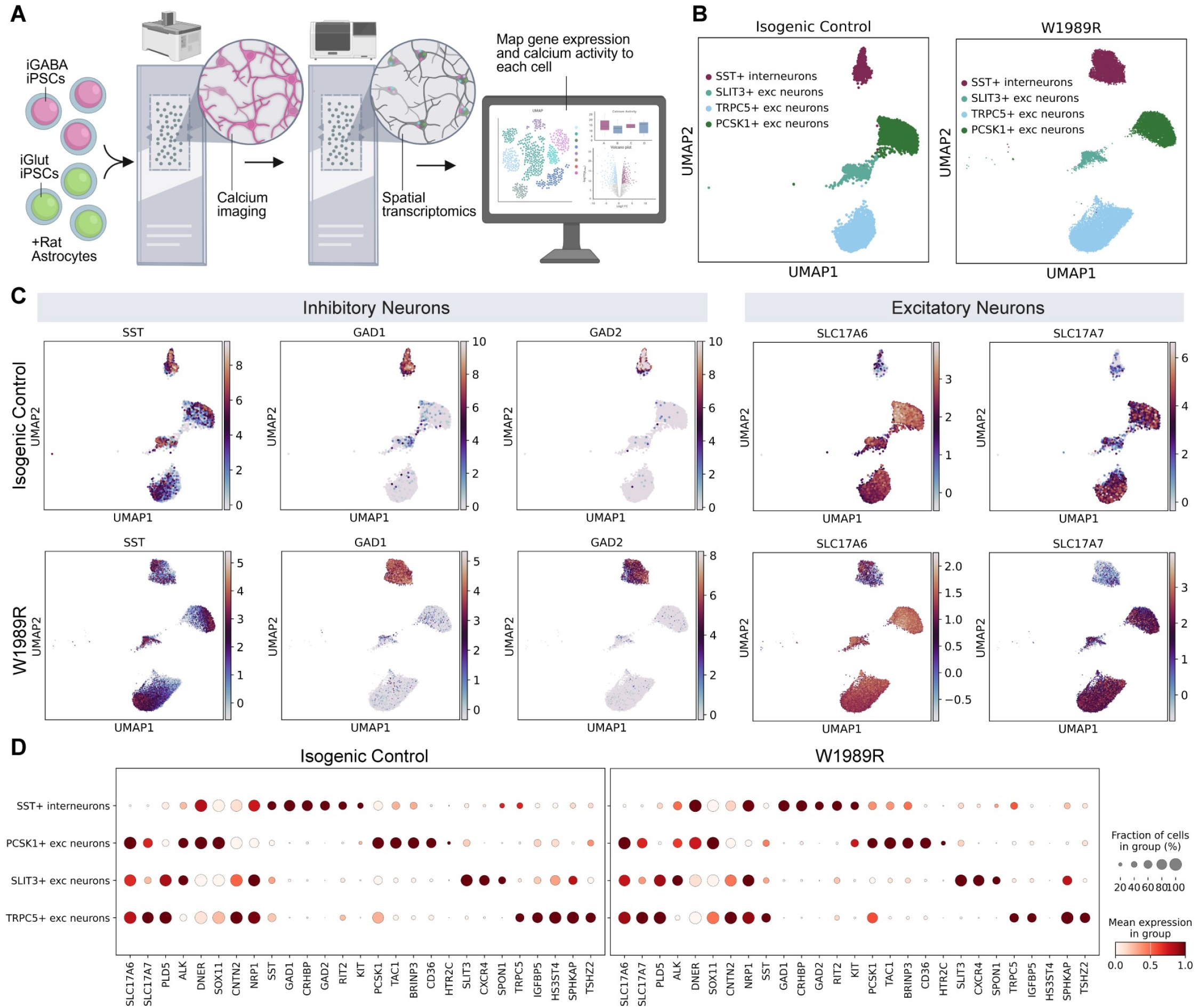
Using VISTA-FISH to measure gene expression and calcium activity in BD patient iPSC-derived neurons. **A:** Diagram of VISTA-FISH approach. Cells are cultured in a custom PDMS gasket on a Xenium 10X Genomics glass slide. Cells were plated onto the glass slide on day three of doxycycline-induced differentiation. BD patient (W1989R) and isogenic control iGlut and iGABA neurons were co-cultured with rat astrocytes until day 45. Live-cell imaging was performed using a confocal microscope to measure neuron activity via calcium transients. Spatial transcriptomic measurement of gene expression was detected using a 10X Genomics Xenium Analyzer and the 266-gene Human Brain panel. Image registration was used to link calcium signaling data to gene expression for each cell. Genes were tested per transcriptomic cluster by Wilcoxon rank-sum test and genes with Benjamini-Hochberg adjusted P < 0.05 and absolute log2 fold-change > 0.5 were called significant. **B:** UMAP plots and cell-type clusters were generated from single-cell gene expression data. Each dot represents a cell. **C:** Expression feature plots of marker genes from each UMAP cluster for excitatory neuron and inhibitory interneuron cell-type clustering. **D:** Dot plot of marker gene expression patterns from each cluster.

Spatial gene expression was measured using the 10X Genomics Xenium platform and a targeted 266-gene human brain panel. Neuronal transcript counts were clustered using Scanpy. UMAP visualization identified one inhibitory neuron cluster and three predominant excitatory neuron clusters in each condition (Figure 4B). The inhibitory neuron cluster expressed the canonical marker genes *GAD1, GAD2*, and *SST*, while excitatory neuron clusters expressed *SLC17A6* and *SLC17A7* (Figure 4C). Excitatory neuron populations were further annotated according to the corresponding cluster-enriched markers *PCSK1, SLIT3*, and *TRPC5* (Figure 4D). These neuronal populations were subsequently used for cell-type-specific comparisons of gene expression and calcium activity between *ANK3* p.W1989R and isogenic control clusters.

### Differential gene expression between *ANK3* p.W1989R and isogenic control neuronal populations

We next compared gene expression between *ANK3* p.W1989R and isogenic control neurons within each transcriptomic cluster. Differential expression analysis identified 99 genes in *PCSK1*+ excitatory neurons, including 67 with higher expression in *ANK3* p.W1989R and 32 with higher expression in isogenic control neurons (Figure 5A). In *SLIT3*+ excitatory neurons, 86 genes were differentially expressed, with 52 higher in *ANK3* p.W1989R and 34 higher in isogenic control neurons. In *TRPC5*+ excitatory neurons, 101 genes were differentially expressed, with 62 higher in *ANK3* p.W1989R and 39 higher in isogenic control neurons. Finally, 92 genes were differentially expressed in *SST*+ interneurons, including 70 with higher expression in *ANK3* p.W1989R and 22 with higher expression in isogenic control neurons (Figure 5A).

**Figure 5.**
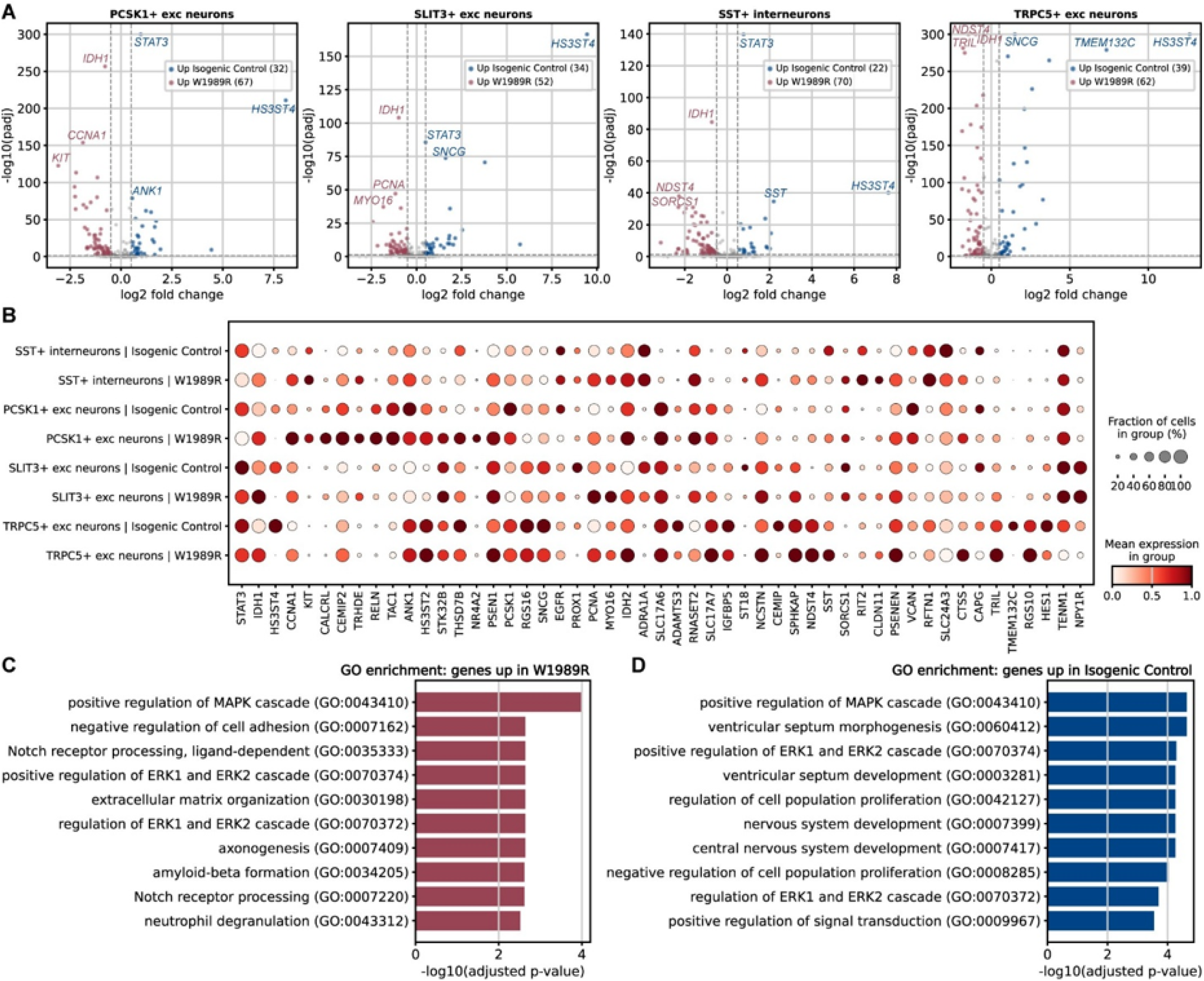
Xenium transcriptomics differential gene expression for *ANK3* p.W1989R versus isogenic control for each neuron type cluster. **A:** Spatial transcriptomic measurement of gene expression was detected using a 10X Genomics Xenium Analyzer and the 266-gene Human Brain panel. Volcano plots showing differential expression between conditions for each cell-type transcriptomic cluster. Genes were tested per transcriptomic cluster by Wilcoxon rank-sum test. Genes with adjusted p < 0.05 (Benjamini-Hochberg) and absolute log2 fold-change > 0.5 were called significant. Significant genes with negative log2 fold-change showed increased expression in W1989R (red), while positive log2 fold-change showed increased expression in isogenic control (blue). **B:** Dot plot of top differentially expressed genes between W1989R and isogenic control for each cluster. **C:** Gene Ontology (GO) enrichment results for genes significantly up in W1989R. **D**: GO enrichment results for genes significantly up in isogenic control.

Several transcriptional differences were shared across neuronal populations (Figure 5B). *IDH1* and *CCNA1*, for example, exhibited higher expression in *ANK3* p.W1989R neurons, while *STAT3* and *HS3ST4* exhibited higher expression in isogenic control neurons. Notably, *SST* expression was lower in the *ANK3* p.W1989R SST+ interneuron cluster than in the corresponding isogenic control cluster.

GO enrichment analysis of genes with higher expression in *ANK3* p.W1989R identified terms related to MAPK and ERK signaling, cell adhesion, extracellular matrix organization, Notch receptor processing, and axonogenesis (Figure 5C). Genes with higher expression in isogenic control neurons were enriched for terms related to MAPK and ERK signaling, nervous system development, cell-population proliferation, and signal transduction (Figure 5D). Because these analyses were restricted to the targeted 266-gene panel, the enriched terms represent candidate biological processes associated with the differential expression of the measured genes rather than an unbiased, transcriptome-wide assessment of gene pathway regulation.

### *ANK3* p.W1989R was associated with cell-type-specific alterations in calcium dynamics

To examine *ANK3* p.W1989R-associated differences in neuronal activity, calcium time-lapse images collected before Xenium analysis were registered to spatial transcriptomic coordinates. Xenium-defined cell boundaries were then used to extract calcium fluorescence traces from individual cells, allowing calcium dynamics to be compared between *ANK3* p.W1989R and isogenic control neurons within each transcriptomic cluster. A multi-tier threshold-based peak-detection algorithm was used to identify discrete calcium transients and quantify calcium transient amplitude, duration, and area under the curve (AUC) (Figure 6).

**Figure 6.**
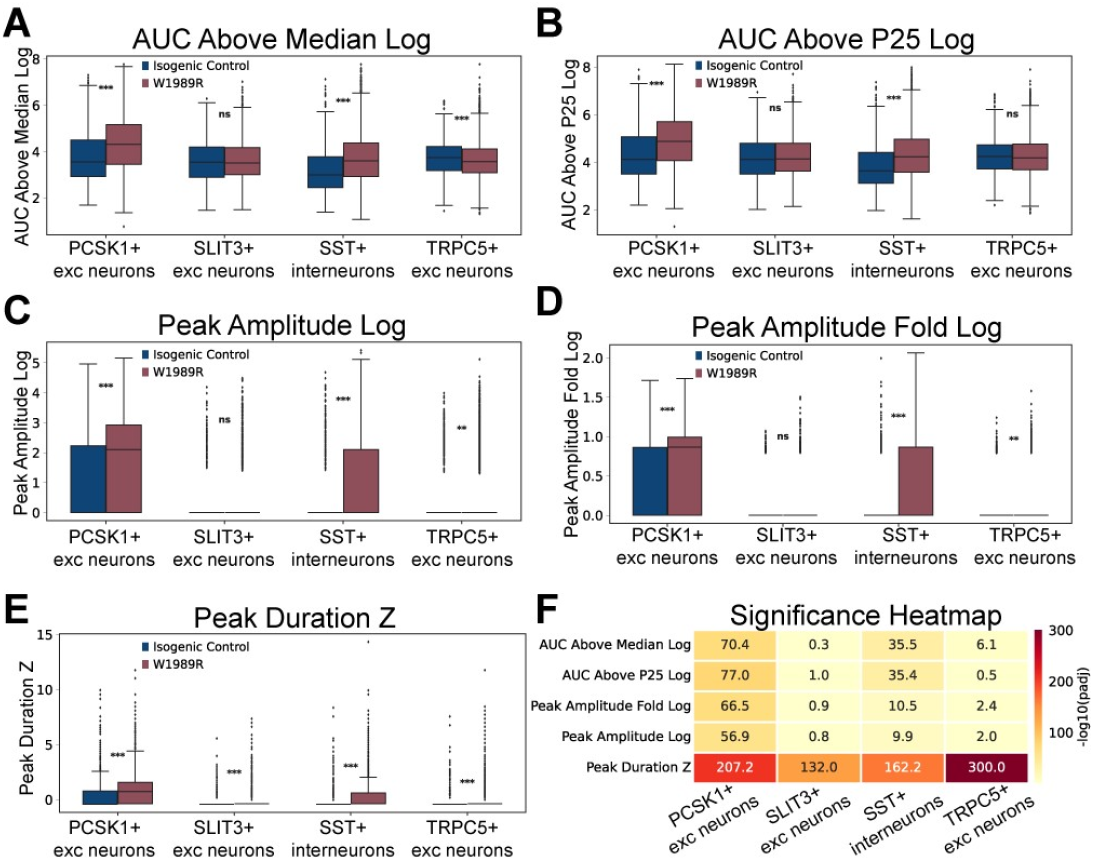
Calcium activity for *ANK3* p.W1989R vs isogenic control for each neuron type cluster. Calcium activity time-lapse fluorescence images from 60 time points were registered to the Xenium transcriptomic coordinate system using an affine transform derived in Xenium Explorer. Mean fluorescence intensity was extracted for each cell by polygon-masking each cell’s Xenium-defined boundary onto the registered image stack. For each cell, a multi-tier threshold-based peak-detection algorithm, scaled to each cell’s own 25th percentile and minimum intensity, identified discrete calcium transients from which we derived: **A:** Area under the curve (AUC) above the median log **B:** AUC above the 25th percentile log **C:** Peak amplitude log **D:** Peak amplitude fold log **E:** Peak duration **F:** Calcium-activity variables were compared between conditions per cluster by Wilcoxon rank-sum test with Benjamini-Hochberg correction across all tests. Comparisons were called significant for padj < 0.05 (−log10 (padj) > 1.3).

AUC above the median was significantly higher in *ANK3* p.W1989R *PCSK1*+ excitatory neurons and *SST*+ interneurons, but lower in *TRPC5*+ excitatory neurons than in the corresponding isogenic control populations (Figure 6A, F). AUC above the 25th-percentile baseline was also higher in *ANK3* p.W1989R *PCSK1*+ excitatory neurons and *SST*+ interneurons, and no significant differences were observed in *SLIT3*+ or *TRPC5*+ excitatory neurons (Figure 6B, F). Peak amplitude measures were significantly higher in *ANK3* p.W1989R *PCSK1*+ excitatory neurons, *SST*+ interneurons, and *TRPC5*+ excitatory neurons, but not *SLIT3*+ excitatory neurons (Figure 6C, D, F). Peak duration was greater in *ANK3* p.W1989R neurons across all four clusters (Figure 6E, F). Collectively, these findings demonstrate cell-type-specific alterations in calcium dynamics in *ANK3* p.W1989R neurons, with *PCSK1*+ excitatory neurons and *SST*+ interneurons showing the most consistent increases in neuronal activity across the measured calcium features.

### SST-mediated cortical inhibition was reduced in heterozygous *ANK3* p.W1989R mice

Altered calcium dynamics and *SST* expression observed in hiPSC-derived *ANK3* p.W1989R *SST*+ interneurons motivated us to investigate how the variant may impact SST interneuron-mediated signaling in the brain using the *Ank3* p.W1989R mouse. Optogenetic stimulation of presynaptic neurons followed by patch-clamp evaluation of postsynaptic cells can be used to study synaptic connectivity and strength in mouse brain slices (46). Cortical SST interneurons preferentially synapse onto pyramidal neuron dendrites to regulate their excitability (47, 48). However, the SST interneuron class is made up of numerous distinct SST interneuron subtypes that vary in their location across cortical layers, their dendritic arbors, and axonal projection patterns (48). These SST interneurons can most easily be divided based on their location in specific cortical layers.

To evaluate if SST synaptic connectivity with pyramidal neurons was altered by *Ank3* p.W1989R, we performed optogenetic stimulation of layer 2/3 and 5/6 SST interneurons in mouse brain slices and recorded the resulting currents from layer 5 pyramidal neurons. SST-Cre^+/-^ x *Ank3* p.W1989R^+/ WT^ or SST-Cre^+/-^ WT mice received prefrontal cortex (PFC) injections of an AAV encoding a Cre-dependent, soma-targeted channelrhodopsin variant (ST-ChroME) (49) (Figure 7A). Paired optical stimuli over 4 light intensities were delivered separately to layers 2/3 and 5/6 to stimulate SST interneurons. The resulting optically evoked inhibitory postsynaptic currents (oIPSCs) were recorded from L5 pyramidal neurons using whole-cell patch-clamp electrophysiology (Figure 7B, C, D).

**Figure 7.**
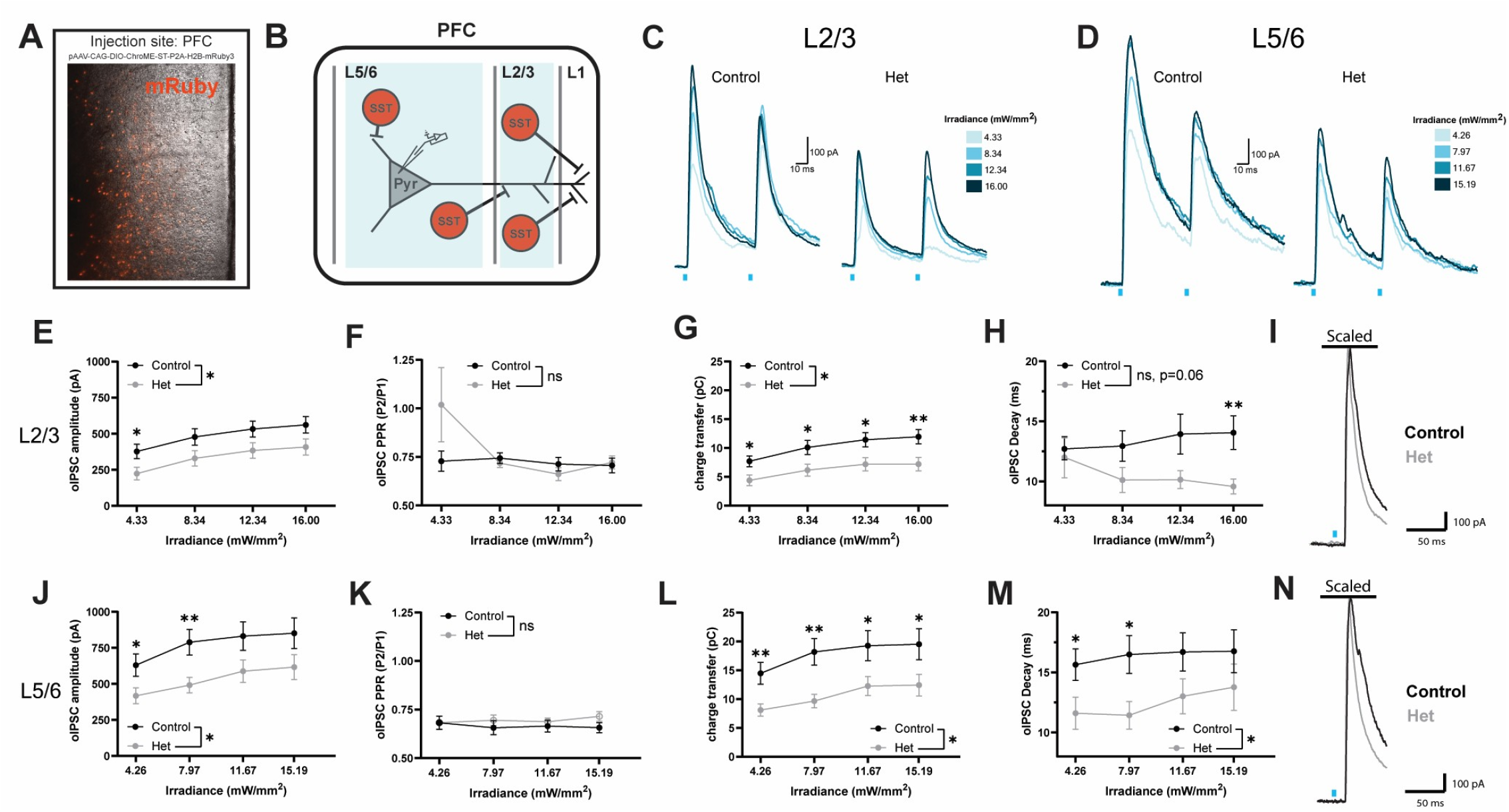
Inhibitory synaptic transmission was altered at SST-L5 pyramidal neuron synapses in the PFC in SST-Cre x *Ank3* p.W1989R mice. **A:** Cre-dependent soma-targeted ChroME was expressed in SST neurons in the PFC of SST-cre mice. **B:** Schematic of selective stimulation of GABA release from L2/3 or L5/6 SST neurons while recording oIPSCs in an L5 pyramidal neuron. The transparent blue area represents the region that optical stimulation was delivered. **C:** Example traces of oIPSCs elicited by optical stimulation of L2/3 SST-INs across four different light intensities in either SST-cre (black) or *Ank3* p.W1989R het (gray) SST-cre mice. **D:** Example traces of oIPSCs elicited by optical stimulation of L5/6 SST-INs across four different light intensities in either SST-cre (black) or *Ank3* p.W1989R het (gray) SST-cre mice. **E:** Summary data of L2/3 oIPSC amplitudes. **F:** Summary data of L2/3 oIPSC charge transfer. **G:** Summary data of L2/3 oIPSC PPR. **H:** Summary data of L2/3 oIPSC decay. **I:** Example traces showing decay of oIPSCs elicited by optical stimulation of L2/3 SST-INs across four different light intensities normalized at pulse 1 peak amplitude in either SST-cre (black) or *Ank3* p.W1989R het (gray) SST-cre mice. **J:** Summary data of L5/6 oIPSC amplitudes **K:** Summary data of L5/6 oIPSC charge transfer. **L:** Summary data of L5/6 oIPSC PPR. **M:** Summary data of L5/6 oIPSC decay. **N:** Example traces showing decay of oIPSCs elicited by optical stimulation of L5/6 SST-INs across four different light intensities normalized at pulse 1 peak amplitude in either SST-cre (black) or *Ank3* p.W1989R het (gray) SST-cre mice.

The peak amplitude of the first oIPSC was significantly lower in slices from SST-Cre^+/-^ x *Ank3* p.W1989R^+/WT^ mice following stimulation of both layer 2/3 and layer 5/6 SST interneurons, suggesting deficits in inhibitory synaptic transmission (Figure 7E, J). The total charge transfer integrated across both the first and second stimuli was also significantly reduced in slices from SST-Cre^+/-^ x *Ank3* p.W1989R^+/WT^ mice following stimulation of both layer 2/3 and layers 5/6 SST interneurons, demonstrating a decrease in the total inhibitory synaptic current over time (Figure 7F, K). Paired-pulse ratio, an indirect measure of presynaptic release probability, did not differ significantly between genotypes following SST interneuron stimulation in either layer, suggesting no alterations in presynaptic GABA release (Figure 7G, L).

The oIPSC decay time to half maximal amplitude, which reflects the duration of the inhibitory current, was significantly shorter in SST-Cre^+/-^ x *Ank3* p.W1989R^+/WT^ mice following stimulation of layer 5/6 SST neurons, while layer 2/3 SST neuron oIPSC half decay time did not reach statistical significance (P = 0.06) (Figure 7H, M). To further characterize altered oIPSC decay kinetics, the first oIPSC of each paired-pulse was fitted with a double exponential decay function, yielding fast and slow decay time constants (τ_fast_ and τ_slow_) (Supplemental Figure 3). τ_fast_ was significantly shorter in SST-Cre^+/-^ x *Ank3* p.W1989R^+/WT^ mice following stimulation of SST interneurons in either layer, while τ_slow_ was unchanged (Supplemental Figure 3A-D). The shorter oIPSC decay time and selective reduction in τfast, with no change in τ_slow_, indicate that the fast component of SST-mediated inhibitory currents decays more rapidly in heterozygous *Ank3* p.W1989R mice. Collectively, these findings identify deficits in SST-mediated inhibition of layer 5 pyramidal neurons in SST-Cre^+/-^ x *Ank3* p.W1989R^+/WT^ mice. Reduced oIPSC amplitude and charge transfer, and faster decay, with unchanged PPR, are consistent with altered postsynaptic GABA_A_ receptor function or organization.

## DISCUSSION

Genome-wide association studies have identified the gene *ANK3*, which encodes ankyrin-G, as one of the most consistently replicated risk loci for bipolar disorder (BD) (10, 11). However, the mechanisms by which *ANK3* BD-associated variants contribute to neuronal dysfunction seen in BD remain poorly understood. Previously, we identified the *ANK3* p.W1989R variant in a family with BD and demonstrated that this variant disrupts inhibitory signaling and leads to neuronal hyperexcitability in mice (19). Here, we sought to further characterize the impact of this variant in neurons derived from an individual with BD who was heterozygous for *ANK3* p.W1989R. CRISPR correction of the variant increased somatodendritic inhibitory synapse density on both excitatory and inhibitory human induced pluripotent stem cells (hiPSC)-derived neurons. Integrated spatial transcriptomic and calcium imaging analyses further identified cell-type-specific differences in gene expression and calcium dynamics, including increased calcium activity in *SST*+ interneurons and *PCSK1*+ excitatory neurons, and decreased *SST* expression in *SST*+ interneurons with the variant. Optogenetic stimulation of prefrontal cortical SST interneurons in heterozygous *Ank3* p.W1989R mice revealed weaker and shorter-lasting inhibition of layer 5 pyramidal neurons. Together, these findings, from complementary human and mouse models, demonstrate reduced inhibitory connectivity and identify SST interneuron-mediated signaling as a component of cortical circuitry affected by *ANK3* p.W1989R.

Our results demonstrate that *ANK3* p.W1989R BD patient-derived excitatory and inhibitory neurons have reduced inhibitory synaptic density compared to their CRISPR-corrected isogenic control counterparts. These findings in excitatory human-derived neurons extend previous work from mouse models showing that ankyrin-G interacts with the GABA_A_ receptor associated protein (GABARAP) through W1989 to limit GABA_A_ receptor endocytosis, and disruption of this amino acid through *ANK3* p.W1989R reduces inhibitory synapse density in cortical and hippocampal pyramidal neurons (18, 19). The present study not only demonstrates that correction of the heterozygous *ANK3* p.W1989R variant is sufficient to increase inhibitory synapse density within the bipolar disorder patient genetic background, but also that the mechanisms of ankyrin-G inhibitory synaptic function appear to be conserved between human and mouse models.

The observed reduction in inhibitory synapse density on *ANK3* p.W1989R patient-derived inhibitory neurons broadens our understanding of the effects of the variant beyond excitatory neurons, suggesting that interneuron-to-interneuron connectivity may also be disrupted by *ANK3* p.W1989R. Inhibitory connections between inhibitory neurons allow for the dynamic control of pyramidal neuron excitability and cortical circuit activity through a process known as disinhibition (50). SST interneurons, which synapse onto the dendrites of pyramidal neurons, receive inhibitory input from multiple interneuron populations, including those expressing vasoactive intestinal peptide (VIP) and parvalbumin (PV) (50, 51). SST interneurons also inhibit PV interneurons, which synapse onto the soma and axon of pyramidal neurons (52). Therefore, it is important that future studies identify specifically which inhibitory neuron subtypes are impacted by this ankyrin-G variant, as sustained hyperactivity of interneurons due to reduced inhibitory synapse density could have differential consequences for cortical circuits depending on what types of inhibitory neurons are impacted.

Spatial transcriptomic analysis identified cell-type-specific and shared transcriptional differences between *ANK3* p.W1989R and isogenic control neurons. Reduced *SST* expression was identified in *ANK3* p.W1989R *SST*+ interneurons. SST interneurons are a diverse class of cortical GABAergic neurons that regulate dendritic integration, cortical plasticity, learning, and network oscillations (26, 27, 51). Reduced SST expression and fewer detectable SST-expressing neurons have been reported in several mood disorders, including BD (27). Importantly, lower *SST* transcript abundance observed in *ANK3* p.W1989R likely reflects a reduction in *SST* expression within the identified interneuron population rather than a reduction in the number of SST interneurons. However, the causes of reduced SST expression in BD, as well as in our study, remain unknown.

*STAT3* expression was also lower in *ANK3* p.W1989R neurons, and interestingly, SST has been shown to attenuate leptin-induced STAT3 phosphorylation in the hypothalamus (53). However, it is unclear whether *SST* and *STAT3* expression levels are dependent on each other, and the parallel reductions in expression observed in *ANK3* p.W1989R neurons may not be directly related. However, the relationship between STAT3 and cancer progression paired with the elevated cancer susceptibility seen in patients with BD highlights the importance of investigating a potential relationship between changes in *SST* and *STAT3* expression in this model (54, 55). Direct measurements of STAT3 phosphorylation and nuclear localization could provide further insight into whether STAT3 phosphorylation or activity is altered in *ANK3* p.W1989R neurons.

Calcium imaging identified cell-type-specific alterations in neuronal activity associated with *ANK3* p.W1989R. *SST*+ interneurons and *PCSK1*+ excitatory neurons exhibited the most consistent increases in calcium-transient amplitude, duration, and integrated activity, while other excitatory populations showed mixed or nonsignificant differences. These findings suggest that functional consequences associated with *ANK3* p.W1989R vary among neuronal populations. These results are consistent with the idea that the variant differentially affects different cell-types, which has been observed in the *Ank3* p.W1989R mouse model (19). It is unknown whether different transcriptomic clusters represent different developmental states of excitatory neurons, however, differences in neuronal maturity could also influence how the *ANK3* p.W1989R variant impacts neuronal gene expression and activity. For *SST*+ interneurons and *PCSK1*+ excitatory neurons, increased calcium activity suggests hyperexcitability consistent with the observed reduction in inhibitory synaptic density. Future studies using patch-clamp electrophysiology on *ANK3* p.W1989R human-derived neuron cultures could be used to characterize the effect of the variant on inhibitory neuron function and further elucidate if deficits in inhibitory signaling seen in excitatory neurons mirror postsynaptic deficits seen in mouse models.

Altered calcium activity and *SST* expression seen in *ANK3* p.W1989R SST+ interneurons motivated us to directly analyze how SST-mediated inhibition may be impacted in the *Ank3* p.W1989R mouse prefrontal cortex. Optogenetic stimulation of layer 2/3 or 5/6 SST interneurons produced smaller oIPSC amplitudes and reduced charge transfer in layer 5 *Ank3* p.W1989R pyramidal neurons, indicating weaker overall SST-mediated inhibition. Paired-pulse ratio was unchanged, providing no evidence for altered presynaptic release probability. The oIPSC decay in heterozygous *Ank3* p.W1989R mice was also faster, with a selective reduction in τ_fast_ and no change in τ_slow_, indicating shorter-lasting inhibition. These results are consistent with altered postsynaptic GABA_A_ receptor organization or kinetics and complement the reduced somatodendritic inhibitory synapse density observed in human *ANK3* p.W1989R neurons. However, it remains unknown what the impact of the *ANK3* p.W1989R variant is on SST interneuron excitability in the cortex, or if other types of inhibitory neurons are impacted by this variant. Future studies investigating which neuron types are impacted in what brain regions will not only provide additional understanding of the impact of the *ANK3* p.W1989R variant but will also expand our knowledge of ankyrin-G-mediated inhibitory synaptic functions.

## MATERIALS AND METHODS

### hiPSC generation

Fibroblasts were obtained and reprogrammed to human induced pluripotent stem cells (hiPSCs) through the Heinz C. Prechter Bipolar Research Program at the University of Michigan. Specifically, fibroblasts were obtained via skin biopsy from the 42-year-old male proband diagnosed with BD I, heterozygous for the *ANK3* p.W1989R variant (19). Fibroblasts were reprogrammed into hiPSCs by the Heinz C. Prechter Bipolar Research Program using the Epi5™ Episomal iPSC Reprogramming Kit (Thermo Scientific, A15960). This was achieved through forced expression of the reprogramming factors Oct4, Sox2, Klf4, and c-Myc (OSKM) used as previously described (33), and as first described as part of the Yamanaka factors (34). Original hiPSC characterization was performed by the Heinz C. Prechter Bipolar Research Program. These hiPSCs are accessible through the New York Stem Cell Foundation (NYSCF) (NYSCF ID: DW0009) and have passed all NYSCF quality control tests for viability, sterility, karyotyping, pluripotency, absence of transgenes, and differentiation capacity. With the help of the Human Stem Cell and Gene Editing (HSCGE) core at the University of Michigan, we generated an isogenic control hiPSC line from these hiPSCs through CRISPR-mediated correction of W1989R. Resulting hiPSCs were sequenced to ensure genetic editing, immunostained for pluripotency markers, and tested to be mycoplasma free through the HSCGE core.

### hiPSC culture

hiPSCs were maintained in mTeSR Plus media (STEMCELL Technologies, 100-0276) with penicillin-streptomycin (Thermo Scientific, 15140122) on plates (Corning, 3516) coated with Geltrex (Thermo Scientific, A1413301) in DMEM/F12 (Thermo Scientific, 11320033) according to the manufacturer’s instructions. At 80% confluency, cells were passed with 1 mM EDTA as described (35) onto new Geltrex-coated plates in mTeSR Plus media (STEMCELL Technologies, 100-0276) with penicillin-streptomycin (Thermo Scientific, 15140122) supplemented with 1 µM Y-27632 (ROCK inhibitor) (Tocris, 1254). The mTeSR Plus media without ROCK inhibitor was replaced within 24 hours, and every 1-2 days after. Mycoplasma testing was performed routinely using the MycoStrip Mycoplasma Detection Kit (InvivoGen, rep-mysnc-100).

### Generation of transgenic doxycycline-inducible hiPSC lines using piggyBac

Stable hiPSC lines expressing doxycycline-inducible transcription factors for neuron differentiation were generated using piggyBac transposition (36). 80% confluent hiPSCs were passed 1:2 into mTeSR Plus (STEMCELL Technologies, 100-0276) with 1µM ROCK inhibitor (Tocris, 1254). The following day, hiPSCs were transfected in E8 media using 2.5 µg of total DNA: To generate induced glutamatergic (iGlut) hiPSCs, cells were transfected with 0.75 µg of pCMV-PBase, 0.25 µg of MT-US2-puro, 0.75 µg of PB-CAG-TetOn-IRES-mCherry and 0.75 µg of PB-TRE-hNGN2. To generate induced GABAergic (iGABA) hiPSCs, cells were transfected with 0.75 µg of pCMV-PBase, 0.25 µg of MT-US2-puro, 0.75 µg of PB-CAG-TetOn-IRES-mCherry and 0.75 µg of PB-TRE-ASCL1-IRES-DLX2. Plasmids were obtained from the Uhler laboratory at the University of Michigan and identity was verified by sequencing. Transfection complexes were made in Opti-MEM (Gibco, 31985-070) according to the Lipofectamine Stem Reagent (Thermofisher, STEM00008) manufacturer protocol and incubated on the cells overnight. 72 h after transfection, cells were treated with 0.8 μg/ml puromycin in E8 media for 48 hours. Following puromycin treatment, cells were maintained in fresh E8 media until colonies of approximately 500 cells were picked manually into individual wells of a 24-well plate. Resulting W1989R and isogenic control iGlut-hiPSC and iGABA-hiPSC clones were expanded, and replicate cultures were tested for neuron differentiation following 72h E8 with 1µM dox treatment.

### iNeuron differentiation

In preparation for differentiation, iGlut-(W1989R iGlut Clone 1, isogenic control iGlut Clone 3) and iGABA-(W1989R iGABA Clone 2, isogenic control iGABA Clone 7) hiPSC lines were plated and expanded as described. To begin differentiation, hiPSCs were passed single cell with Accutase (STEMCELL Technologies, 07920) in mTeSR Plus media (STEMCELL Technologies, 100-0276) with 1µM ROCK inhibitor (Tocris, 1254) onto 10 cm Geltrex-coated plates (Corning, 430167) at 3 million cells per plate. Starting on DIV0, cells were maintained in fresh E8 media (STEMCELL Technologies, 05990) with 1µM doxycycline (Sigma-Aldrich, D9891) until DIV3.

In preparation for DIV3 passaging, plates were coated overnight at room temperature with 2.2 mg/mL PEI (Sigma-Aldrich, 408727) in 0.1M Borate Buffer pH 8.4 (Boric acid (Sigma-Aldrich, B7901) RNase, DNase-free water (Thermo Scientific Chemicals, J60610-EQC), sterile filtered through 0.22 μm PES filter system (CELLTREAT, 229707)). The next day, plates were washed with sterile water five times and left open to dry for one hour in a sterile biosafety cabinet. Plates were then coated with laminin (Sigma-Aldrich, L2020-1MG, diluted 1:40) in DMEM/F12 and incubated for at least 1 hr or overnight at 37°C. Plastic six-well plates (Corning, 3516) were used to obtain cell pellets, glass-bottom plates (Cellvis, D35-20-1.5-N) were used for imaging, and Xenium 10X Genomics glass slides were used for VISTA-FISH analysis.

On DIV3, the iGlut- and iGABA-NPCs were plated on PEI- and laminin-coated plates to create excitatory/inhibitory co-cultures at a 50:50 ratio. Cells were passaged as single cells using Accutase (STEMCELL Technologies, 07920) in 3N differentiation media (250 mL DMEM/F-12 (Thermo Scientific, 11320033), 250 mL Neurobasal (Thermo Scientific, 21103049), 125 μL of 10 mg/mL insulin (Sigma I9278), 2.5 mL Non-Essential Amino Acids (Gibco 11140-050), 2.5 mL N2 supplement (Gibco 17502-048), 5mL B27 supplement (Gibco 17504-044), 2 µl β-mercaptoethanol stock (12 M) (Sigma M7522), 2.5 mL GlutaMAX (Gibco 35050-061), 4mL penicillin-streptomycin (Thermo Scientific, 15140122) (37) with 1µM doxycycline and 1µM ROCK inhibitor. Cells were counted using a fluorescence automated cell counter (Invitrogen™, A49866). The next day (DIV4), a full media change to new 3N media was performed with 1µg/mL doxycycline, 1:200 Geltrex, and thawed cryopreserved rat astrocytes (Sartorius, 4586). Astrocytes were added at a 1:1 ratio of the total iGlut and iGABA cell count plated. On DIV5, a half-media change was performed with 3N media containing 1µg/mL doxycycline and AraC (Sigma-Aldrich, C6645). Half-media changes to fresh 3N media with 1µM doxycycline were performed until DIV8, when a full media change was performed with BrainPhys media (BrainPhys media kit (STEMCELL Technologies, 05792)), 0.02 μg/1mL BDNF (PeproTech, 450-02), 0.02 μg/1mL GDNF (PeproTech, 450-10), 5 M HEPES (Thermo Scientific, 15630080), penicillin-streptomycin (Thermo Scientific, 15140122). BrainPhys half-media changes were performed three times a week until analysis. Media was supplemented with Geltrex (1:1000) once a week for the entirety of the differentiation. This differentiation protocol was obtained and adapted from the HSCGE core at the University of Michigan.

### Immunocytochemistry

Pluripotency marker immunocytochemistry for Nanog (Abcam, Ab21624), Oct3/4 (Abcam, Ab181557), SSEA4 (Abcam, Ab16287), and SOX2 (Millipore, AB5603) was performed on isogenic control hiPSCs by the HSCGE core at the University of Michigan. Cell samples were washed at room temperature in 1X PBS, pH 7.4 (Thermo, 10010023), and then fixed for 15 minutes in 4% paraformaldehyde. Samples were washed three times with PBS (5 minutes each) then incubated for 10 minutes in PBS with 0.2% Triton-X-100. Samples were incubated in blocking buffer (PBS with 0.05% Tween-20, 5% normal goat serum, and 1% BSA) for one hour at room temperature. Samples were then incubated with primary antibody diluted according to the manufacturer’s instructions in blocking buffer overnight at 4 ºC. Samples were washed three times (10 minutes each) with PBS with 0.05% Tween-20. Samples were then incubated with secondary antibody (1:400) in blocking buffer at room temperature for 60-120 minutes. Samples were washed 3 times (five minutes each) in PBS with 0.05% Tween-20, then incubated for five minutes in PBS with bis-benzimide (Hoechst stain) to label nuclei. Samples were washed three times (10 minutes each) in PBS with 0.05% Tween-20. Samples were mounted in warm Glycergel and allowed to cure overnight before imaging. Immunostaining protocol provided by the HSCGE core at the University of Michigan.

To evaluate the number of inhibitory synapses and axon initial segments (AIS), immunocytochemistry was performed on iGlut/iGABA cultures with rat astrocytes on DIV45 of neuronal differentiation. The following primary antibodies were used: rabbit anti-ankyrin-G (1:1000, Jenkins Laboratory (38)), mouse anti-MAP2 (1:1000, Millipore, MAB3418), and guinea pig anti-vGAT (1:1000, Synaptic Systems, 131-004). Secondary antibodies used were Alexa Fluor 488 donkey anti-rabbit IgG (1:250, Thermo Scientific, A21206), Alexa Fluor 405 donkey anti-mouse IgG (1:250, Thermo Scientific, A48257), and Alexa Fluor 568 goat anti-guinea pig IgG (1:250, Thermo Scientific, A11075). To identify green excitatory neurons versus red inhibitory neurons, nanobodies FluoTag®-X4 anti-GFP conjugated to Atto488 (1:500, NanoTag, N0304) and FluoTag®-X4 anti-RFP conjugated to Atto565 (1:500, NanoTag, N0404) were used. An anti-gephyrin nanobody conjugated to SyliteD1.1-SC5 (1:250, NanoTag, P4001) was also used. Samples were fixed in 4% paraformaldehyde (PFA) (16% PFA (Thermo Scientific, 28906) diluted in 1XPBS pH 7.4 (Thermo Scientific, 10010049)) for 15 minutes at room temperature, followed by 0.2% Triton (Sigma-Aldrich, T8787) in 1XPBS pH 7.4 for 10 minutes. Blocking buffer (5% bovine serum albumin (GeminiBio, 700-100P) in PBST (0.2% TWEEN 20 (Sigma-Aldrich, P1379) in 1XPBS pH 7.4)) was added for 30 minutes. Primary antibodies were diluted in blocking buffer and added to samples overnight at 4°C. The next day, samples were washed with PBST three times for five minutes each. Secondary antibodies and nanobodies were diluted in blocking buffer on ice and added to the samples following the final PBST wash. The covered samples were incubated at room temperature for one hour, and the three, five minute, PBST washes were repeated. Samples were then post-fixed in 4% PFA for 30 minutes at room temperature. Cells were then washed twice with PBST for five minutes. Following the final wash, cells were mounted in ProLong® Gold Antifade Reagent (Thermo Scientific, P36930).

### Imaging

Confocal images of immunocytochemistry for inhibitory synapses and AIS properties in DIV45 samples were acquired using a Zeiss LSM 880 with Airyscan using a 63x 1.4 Plan-Apochromat objective. The 405, 488, 561, and 633 nm lasers were used for excitation, and laser power and imaging parameters were kept constant for each experimental condition and sample. All image acquisition for experimental conditions was performed blinded.

### Inhibitory synapse quantification

Z-stack confocal CZIs were processed using Airyscan processing. CZIs were converted to Imaris images using Imaris software. Methods were adapted to create 3D somatodendritic surfaces from each neuron using the MAP2 signal, which was then used to identify inhibitory synapses (39). Inhibitory synapses were quantified as the number of postsynaptic gephyrin puncta with a diameter of 0.3µm within the MAP2 surface colocalized within 1 µm of presynaptic vGAT puncta with a diameter of 0.3µm outside the MAP2 surface. MAP2 surface volume was calculated and used to normalize the number of synaptic gephyrin puncta to the size of the MAP2 surface. All image analysis was performed blinded.

### Axon initial segment (AIS) quantification

Z-stack confocal CZI files were processed using Airyscan processing, then used to create maximum intensity projections. ImageJ (Fiji) was used to draw regions of interest (ROIs) starting at the soma, and down past the end of the AIS. The soma and AIS were identified by MAP2 and ankyrin-G staining, respectively. Intensity profiles of ankyrin-G fluorescence were generated for each ROI. A background fluorescence intensity measurement was also taken in the same channel. Background fluorescence was subtracted from the ankyrin-G fluorescence intensity plot profile. The start and end of the AIS were defined as the points where ankyrin-G fluorescence intensity reached 15% of the maximum fluorescence intensity for each ROI. For neurons with multiple AISs, each AIS was measured and averaged per neuron.

### Immunocytochemistry statistics

Statistical analyses were performed using GraphPad Prism (GraphPad Software). Detailed statistical comparisons and descriptions are reported in each respective figure legend. For all comparisons, n (number of cells) and N (number of differentiation replicates) are both reported.

### Spatial transcriptomics and cell-type annotation

Xenium in situ spatial transcriptomics (10x Genomics, 266-gene targeted human brain panel) was performed on DIV45 isogenic control iGlut/iGABA neurons and W1989R patient-derived iGlut/iGABA neurons, co-cultured with rat astrocytes. Raw transcript counts were filtered to cells with >=100 total counts, normalized (total-count normalization, log1p), and clustered (PCA, k-nearest-neighbor graph, Leiden clustering) using Scanpy. Clusters were assigned cell-type identities from canonical marker genes (e.g., *GAD1*/*GAD2*/*SST* for interneurons; *SLC17A6*/ *SLC17A7* and cluster-specific markers *SLIT3, TRPC5*, and *PCSK1* for excitatory neuron subtypes; *GJA1*/*SOX9* for astrocytes) and visualized by UMAP. For the neuron-focused reanalysis presented here, both astrocyte clusters were removed from each sample, and in W1989R a minor *NWD2*+ excitatory cluster was merged into the transcriptionally similar *TRPC5*+ cluster; gene-expression normalization and differential expression were then rerun on this neuron-only cell population. Cell-type-specific marker expression was visualized with dot plots (mean expression, scaled per gene) and single-gene UMAP overlays. Physical (x,y) cell centroids from the Xenium output were used for spatial plots of cluster identity.

### Calcium imaging and feature extraction

Calcium activity time-lapse fluorescence images (60 time points) were registered to the Xenium coordinate system using an affine transform derived in Xenium Explorer, and per-cell mean fluorescence intensity was extracted by polygon-masking each cell’s Xenium-defined boundary onto the registered image stack, with pixel intensities clipped at 500 to limit outlier influence. For each cell, a multi-tier threshold-based peak-detection algorithm (thresholds scaled to each cell’s own 25th-percentile and minimum intensity) identified discrete calcium transients, from which we derived peak count, peak amplitude (absolute and fold-change), peak duration, and area under the curve above the median/25th-percentile/minimum baseline. Because raw fluorescence scales differ between imaging sessions, W1989R intensities were treated as the reference distribution and isogenic control intensities were quantile-normalized (rank-based percentile mapping) onto it prior to recomputing all calcium features, so that downstream comparisons and clustering used values on a common scale.

### VISTA-FISH Statistics

Differential expression between W1989R and isogenic control, and marker genes used to identify cell-type clusters within each condition, were tested per transcriptomic cluster by Wilcoxon rank-sum test (Scanpy rank_genes_groups); genes with adjusted P < 0.05 (Benjamini-Hochberg) and absolute log2 fold-change > 0.5 were called significant. Calcium-activity variables were compared between conditions per cluster by Wilcoxon rank-sum test with Benjamini-Hochberg correction across all tests. Gene Ontology enrichment (Biological Process and Molecular Function, Enrichr 2021 libraries) was performed on significant differentially expressed genes using gseapy, with the background gene-universe size set to the 266-gene Xenium panel rather than the whole genome, reflecting the targeted nature of the assay.

### Animals

SST-IRES-Cre (SST-Cre) mice (Source: The Jackson Laboratory, strain #: 018973) were backcrossed onto C57/BL6J, *ANK3* p.W1989R mice were described previously (19). SST-IRES-Cre heterozygous and *ANK3* p.W1989R heterozygous male and female mice were crossed to generate respective genotypes used for all experiments. Animals were group-housed (two to five per ventilated cage) with 24-hour (12/12 dark/light cycle) access to food and water. Litters from the breeders were weaned at postnatal day 21 (P21) and raised to P60-120. All experimental procedures were performed in accordance with the guidelines for animal care of the Institutional Animal Care and Use Committee (IACUC), and University Laboratory Animal Management (ULAM) at the University of Michigan.

### Stereotaxic Injections

Stereotaxic injections were performed as previously described (40) to deliver a Cre-dependent soma-targeted channelrhodopsin variant (ChroME). Mice were injected bilaterally with an AAV Type 9 encoding ChroME [AAV9-CAG-DIO-ChroME-ST-P2A-H2B-mRuby3] targeting the PFC (Addgene viral prep #108912, gift from Hillel Adesnik). In total, approximately 100 nl of virus was injected into the PFC (A/P, +1.7 mm; M/L, ±0.3 mm; D/V, 2.35–2.45 mm).

### Brain Slice Electrophysiology

Brain slice preparations were performed as previously described (41). The 300 µm coronal sections containing the PFC (bregma: 1.94 mm – 1.34 mm) were made and recovered in warmed carbogenated Krebs’ solution supplemented with 10 µM MK-801 (33°C for 30 min). Slices were then maintained in carbogenated Kreb’s solution at room temperature. For PFC recordings, borosilicate glass patch pipettes (Sutter Instrument) were pulled to a resistance of 2.0–4.0 MΩ and filled with a cesium gluconate-low chloride–based internal solution (in mM: 135 cesium gluconate, 1 EGTA, 1.5 MgCl2, 10 HEPES (Na), 2 Na ATP, 0.4 Mg GTP, 7.8 Na2 phosphocreatine). Slices were placed in the recording chamber and continuously perfused with carbogenated Krebs’ solution at 32–34°C. Layer 5 pyramidal (L5 PYR) neurons were identified based on cell morphology. Whole-cell recordings were made with a MultiClamp 700B amplifier (Molecular Devices) digitized at 20 kHz (National Instruments BNC-2090A). Data acquisition was performed using MATLAB-based (MathWorks) WaveSurfer software (HHMI Janelia, RRID:SCR_021529, https://wavesurfer.janelia.org/). Series resistance was monitored throughout the recordings, and only recordings in which the series resistance remained <12 MΩ and did not increase >25% were included. To isolate optically evoked inhibitory postsynaptic currents (oIPSCs), voltage–clamp recordings were made from L5 pyramidal neurons at +5 mV holding potential (not corrected for liquid junction potential). Cells were dialyzed for 5 min before experimental recordings.

Layer-specific oIPSCs were elicited every 30 s by LED illumination (470nm) through the microscope objective (20x Olympus BX51WI) using a digital mirror-device based patterned illuminator (Polygon 400, Mightex). Rectangular ROIs were created to cover L2/3 and L5/6 of the PFC to deliver local optical stimulation to each area selectively. ROI dimensions and position were kept constant between all experiments. Paired optical stimuli were generated by delivering two LED pulses at a 50 ms interval (1ms width, 20 Hz). 4 optical stimuli of increasing irradiance were delivered to each ROI (Layer 2/3: 4.33, 8.34, 12.34, and 16.00 mW/mm2/ Layer 5/6: 4.26, 7.97, 11.67, and 15.19 mW/mm2). Input-output curves were generated by recording and averaging oIPSCs from 5 sweeps following stimulation across each ROI and across each irradiance.

### Analysis of Electrophysiology Data

Raw data were analyzed with Matlab using custom scripts. Sweeps were baseline-subtracted, filtered with a 1 kHz low-pass Gaussian filter, averaged and plotted. oIPSC amplitude of pulse 1 was calculated as the maximum value to its relative baseline. Paired-pulse ratio (PPR) was calculated by dividing the peak amplitude of pulse 2 by the peak amplitude of pulse 1. Incomplete decay of the P1 oIPSC was accounted for by baseline correcting the P2 oIPSC. The P2 baseline was estimated by fitting a double exponential decay function to the P1 decay and extrapolating that decay to the time of the P2 peak. The residual P1 current amplitude at the time of the P2 peak was used as the baseline to estimate the corrected P2 amplitude. Charge transfer was calculated by using the trapezoidal rule formula to estimate the total integrated area under both P1 and P2 oIPSC waveforms. oIPSC decay time was measured as the time it takes for the oIPSC to decay from its peak to 50% of its maximum amplitude.

### Statistics of Electrophysiology Data

Statistical analyses were performed using GraphPad Prism (GraphPad Software). Detailed statistical comparisons and descriptions are reported in each respective figure legend.

## ABBREVIATIONS USED

BD: Bipolar Disorder
GWAS: Genome-Wide Association Studies
GABARAP: GABA_A_ Receptor-Associated Protein
hiPSC: Human Induced Pluripotent Stem Cell
SST: Somatostatin
oIPSC: Optically Evoked Inhibitory Postsynaptic Current
AIS: Axon Initial Segment
GABA: Gamma-Aminobutyric Acid
LIR: LC3-Interating Region
CRISPR: Clustered Regularly Interspaced Short Palindromic Repeats
IC: Isogenic Control
AUC: Area Under the Curve
PFC: Prefrontal Cortex
WT: Wild Type

## ACKNOWLEDGEMENTS

We thank the University of Michigan Heinz C. Prechter Bipolar Research Program for providing the hiPSCs used in this study, and the University of Michigan Human Stem Cell and Gene Editing Core for their assistance with hiPSC genetic editing. We also thank Dr. Lori Isom, Dr. Kristen Verhey, Dr. Sami Barmada, and Dr. Michael Uhler for their valuable feedback on this manuscript.

## AUTHOR CONTRIBUTIONS

KPD and PMJ conceived the project. KPD, SCV, CCK, CWS, LYM, JSS, CJD, MY, KHL, and JMR performed experiments and analyzed data. KPD wrote the original draft of the manuscript with contributions from SCV, MY, and JMR. All authors participated in the editing and final design of the manuscript.

## SUPPLEMENTARY MATERIALS

Figures S1 to S3

