## Supplemental Figures for "Human and Mouse Models of the Bipolar Disorder-Associated *ANK3* p.W1989R Variant Display Altered Somatostatin Interneuron Connectivity and Function"

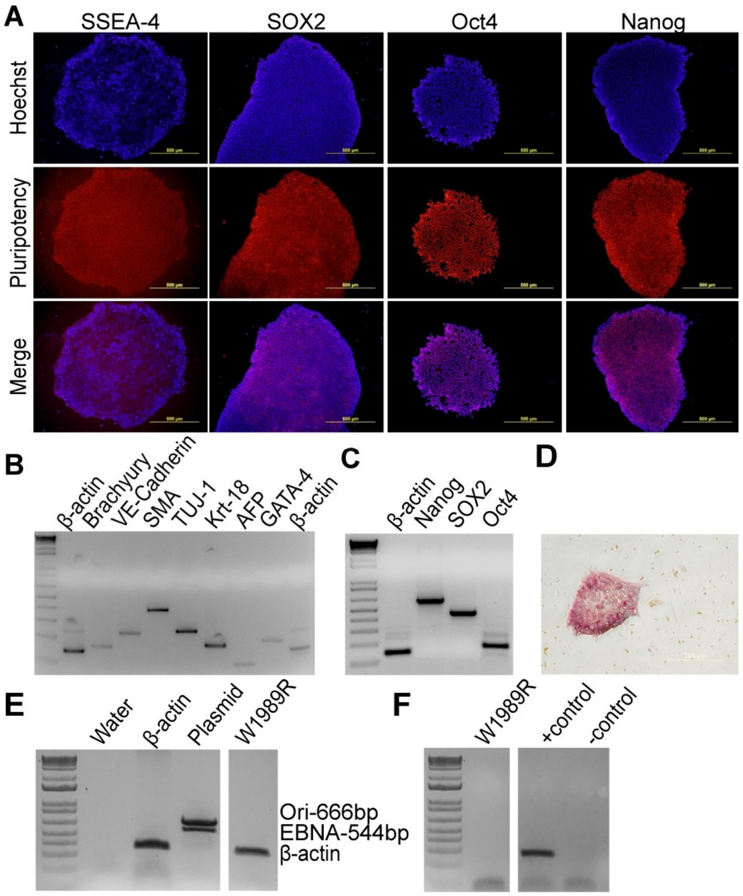


**Supplemental Figure 1. Generation and characterization of human induced pluripotent stem cells (hiPSCs) from a BD patient with the *ANK3* p.W1989R variant.**

**A.** Immunocytochemistry of pluripotency markers SSEA-4, SOX2, Oct4, and Nanog. DNA is Hoechst stained (blue). **B.** Germ layer marker PCR for Brachyury, VE-cadherin, SMA, TUJ-1, Krt-18, AFP, and GATA-4 following embryoid body formation with β-actin control. **C.** Pluripotency marker PCR for Nanog, SOX2, and Oct4 with β-actin control. D. Alkaline phosphatase (AP) pluripotency staining. **E.** Episomal vector PCR test. **F.** Mycoplasma PCR test. Reprogramming and characterization performed by the Heinz C. Prechter Bipolar Research Program and the HSCGE core at the University of Michigan.

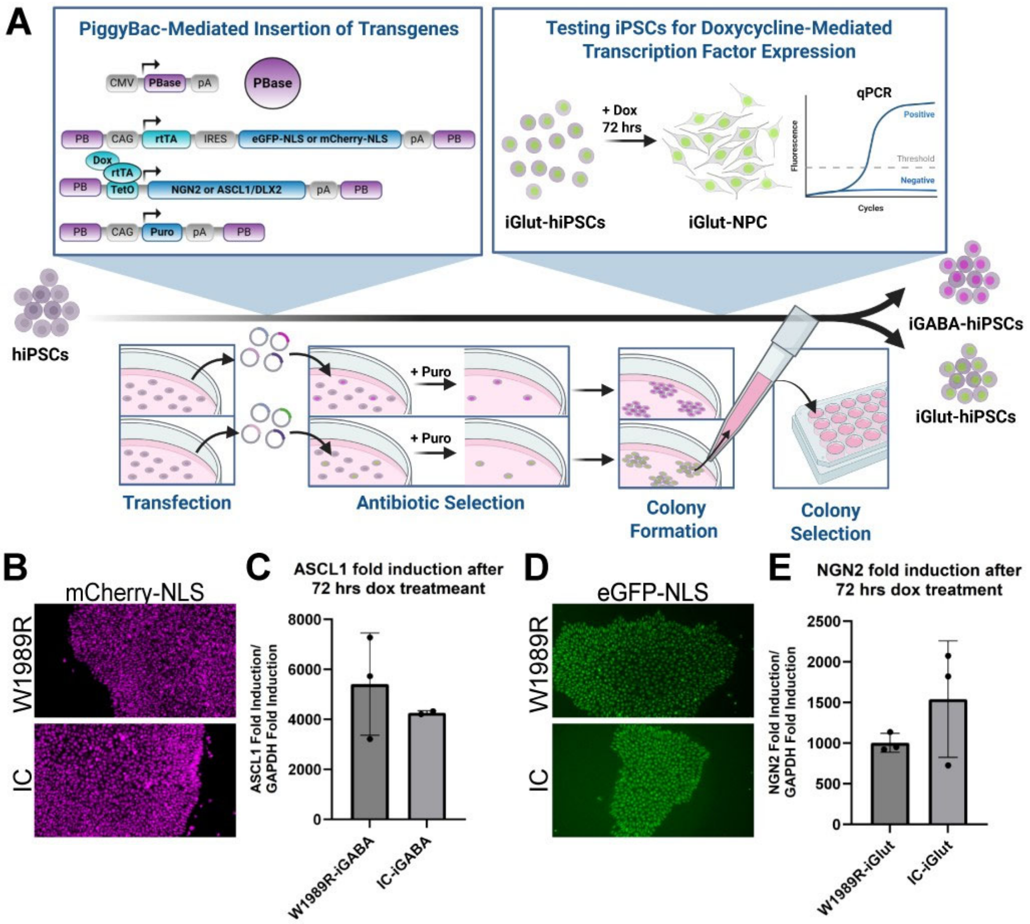


**Supplemental Figure 2. PiggyBac-mediated insertion of doxycycline-inducible transcription factors into hiPSCs.**

**A.** Schematic for methods used to generate doxycycline-inducible GABAergic (iGABA-) and glutamatergic (iGlut-) hiPSCs. PiggyBac-mediated insertion of transgenes encoding puromycin resistance (Puro), nuclear fluorescent marker (eGFP-NLS or mCherry-NLS), and doxycycline-inducible (rtTA, TetO) transcription factors (NGN2 or ASCL1/DLX2). iGlut-hiPSCs: eGFP-NLS and NGN2. iGABA-hiPSCs: mCherry-NLS and ASCL1/DLX2. Colonies were selected for puromycin resistance and nuclear colony fluorescence. iGlut- and iGABA-hiPSCs were treated with doxycycline for 72 hrs, with successful clones displaying neural progenitor cell (NPC)-like morphology and corresponding transcription factor induction analyzed by RT-qPCR. **B.** iGABA-hiPSC W1989R and isogenic control clones display nuclear Cherry-NLS (magenta). **C.** iGABA-hiPSC W1989R and isogenic control clones display ASCL1 induction by RT-qPCR after 72 hours of doxycycline treatment. **D.** iGlut-hiPSC W1989R and isogenic control clones display nuclear eGFP-NLS (green). **E.** iGlut-hiPSC W1989R and isogenic control clones display NGN2 induction by RT-qPCR after 72 hours of doxycycline treatment. For RT-qPCR data, n = 3, N = 2-3.

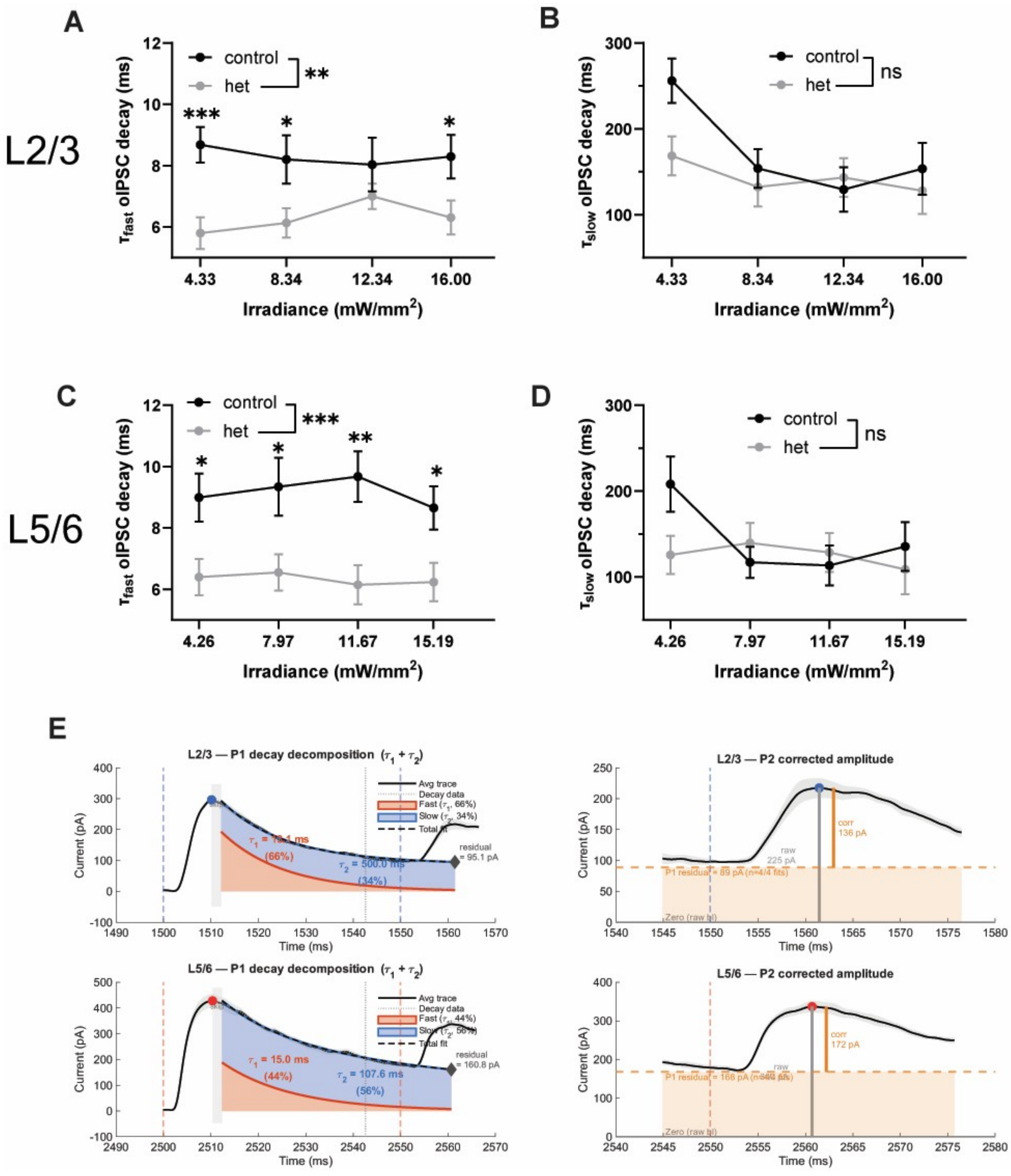


**Supplemental Figure 3. The fast oIPSC decay time constant was selectively altered in SST-Cre x Ank3 p.W1989R mice regardless of layer.**

**A**. Summary data of L2/3 $\tau_{fast}$ oIPSC decay. **B**. Summary data of L2/3 $\tau_{slow}$ oIPSC decay. **C.** Summary data of L5/6 $\tau_{fast}$ oIPSC decay. **D.** Summary data of L5/6 $\tau_{slow}$ oIPSC decay. **E**. Example curve fitting of pulse 1 oIPSC decay and pulse 2 baseline extrapolation.
